# A unified evolutionary framework to reconstruct the viral histories of human, simian, and prosimian immunodeficiency viruses across timescales

**DOI:** 10.1101/2023.06.28.546833

**Authors:** Mahan Ghafari, Peter Simmonds, Aris Katzourakis

**Affiliations:** Department of Biology, Peter Medawar Building for Pathogen Research, University of Oxford, Oxford, UK; Nuffield Department of Medicine, Peter Medawar Building for Pathogen Research, University of Oxford, Oxford, UK

## Abstract

The evolutionary history of simian and prosimian immunodeficiency viruses (SIVs and pSIVs) remains difficult to resolve due to time-dependent rate variation in viral evolution. Standard molecular clock methods relying on contemporary sequences often underestimate viral ages, conflicting with paleovirological and biogeographical evidence. Here, we apply the Prisoner of War model, which accounts for time-dependent rate effects, to reconstruct the evolutionary history of primate lentiviruses. Using non-recombinant genomic regions from 436 HIV, SIV, and pSIV sequences, we estimate that the most recent common ancestor of extant SIVs dates to ∼1 million years ago. The lineages giving rise to HIV-1 and HIV-2 (SIVcpz and SIVsmm) have circulated in their respective hosts for tens to hundreds of thousands of years, suggesting prolonged pre-pandemic human exposure. The divergence of SIV and pSIV is dated to 17–51 million years ago, consistent with lentiviral transfer during the last mammalian colonisation of Madagascar. Our analysis integrates lentiviral data across recent (HIV), intermediate (Bioko Island SIVs), and ancient (pSIV) timescales, enabling inferences not possible from any single timescale alone, and recovers both shallow and deep divergence events without relying on fixed internal node calibrations. This unified framework provides new insights into virus–host coevolution and zoonotic risk.

## Introduction

The discovery of the human immunodeficiency viruses types 1 and 2 (HIV-1, HIV-2) in the 1980s [1–3] led to the identification of a larger and diverse group of primate lentivirus known as simian immunodeficiency viruses (SIVs). These close relatives of HIV are distributed in various non-human apes and Old World monkey species in sub-Saharan Africa including chimpanzees, gorillas, and sooty mangabeys. Subsequent to their discovery, there were extensive investigations into the origins of HIV and identification of potential zoonotic transmission events from SIVs to humans [4]. These studies had revealed unequivocally that SIVcpz transmissions from chimpanzee subspecies, Pan troglodytes troglodytes, and SIVsmm transmission from sooty mangabeys, Cercocebus torquatus atys, were responsible for the emergence of most or all of the HIV-1 and HIV-2 strains respectively in humans [5–7].

Molecular epidemiological studies based on extant SIV and HIV sequences showed a very high rate of nucleotide sequence change over time that predicted a very recent evolutionary origin for primate lentiviruses [4,8]. Using a molecular clock rate estimate of around 10^-3^ substitutions per site per year (s/s/y) for HIV and extrapolation of that across the entire primate lentivirus phylogeny predicts that the most recent common ancestor (tMRCA) of the entire group might have lived no earlier than a couple of centuries ago. More advanced evolutionary reconstructions based on relaxed clocks which, to a certain degree, account for rate variation across different taxa, would push the common ancestor of extant SIV sequences in chimpanzee back to a few hundred to 2,000 years ago [9]. However, all of these estimates based on molecular clock analysis stand in stark contrast to palaeobiological evidence from endogenous viral elements (EVEs) that show lentiviruses have infiltrated their hosts’ germlines over millions of years. These examples include the rabbit endogenous lentivirus type K which became endogenised in European rabbits about 12 million years ago and two prosimian endogenous lentiviruses, fat-tailed dwarf lemur prosimian immunodeficiency virus (pSIVdfl) and grey mouse lemur prosimian immunodeficiency virus (pSIVgml), that independently integrated into the lemur’s genome nearly 4 million years ago [10,11]. The remarkable genetic similarity of these integrated virus elements to currently circulating lentiviruses suggests that exogenous viruses have evolved at an apparently much slower rate over deeper timeframes than predicted by standard molecular clock models.

Molecular clock predictions of a recent evolutionary origin for HIV-1 and HIV-2 are supported by their initially restricted distribution in Central and West Africa - if either had widely infected humans much earlier than the late nineteenth century, they would have been exported to the Americas and northern Europe via slave trade routes between the sixteenth and nineteenth century and likely persisted in those populations until the present. However, predictions of relatively recent origins for the more divergent strains of SIV are contradicted by the results of an analysis of strains of SIV infecting four simian species on the island of Bioko. The fauna of this island has been separated from the mainland for around 10,000 years following the rise in sea levels at the end of the last ice age [12]. Therefore, SIV strains infecting these monkeys must have diverged from their mainland relatives for at least this period. However, SIV strains showed much less sequence divergence than would have been expected by extrapolation of short-term substitution rates. Indeed, using the island separation as a minimum biogeographic calibration point for their divergence, the authors were able to push the tMRCA estimate of SIV strains infecting monkeys and apes in Africa back to nearly 76,000 years ago, with a corresponding clock rate estimate of approximately 10^-5^ s/s/y over this period. However, they also noted that their estimate for the origins of SIV is still likely an underestimate as they found evidence of substitution saturation in their phylogenetic analysis [12].

What the Bioko Island study also highlighted the inability of standard molecular clock methods to account for time-dependent changes in SIV evolutionary rate estimates across different timescales. While the biogeographic calibration point allowed for the estimation of the divergence of some SIV strains infecting monkeys across intermediate timescales (c.a. tens of thousands of years), the same rate could not be used to estimate the short-term evolutionary changes in SIV or HIV because the virus appears to be evolving at a much faster rate over shorter timescales (c.a. hundred years or less). The inferred rate from the biographic calibration point is also not appropriate to estimate the origins of SIV over longer timescales because the substitution rate of the virus is likely even slower as we go further back in time. However, so far, no molecular clock method has been able to explain these changes in the substitution rate estimates for SIV across different timescales under a unified evolutionary framework. Indeed, the existence of a time-dependent evolutionary rate for primate lentiviruses and other organisms [13,14] would create major difficulties in reconstructing virus evolutionary timescales using standard models, creating a substantial underestimation of the timescale for deeper evolutionary histories of a virus if a shallow calibration point is chosen for the analysis. Conversely, the relatively slow rate estimate based on palaeobiological and biographic calibration points, such as from the Bioko study, creates an unrealistically lengthened timescale for the more recent emergence of HIV-1 and other lentiviruses estimated from recent sampling.

The evolutionary history of primate lentiviruses is further complicated by frequent interlineage recombination [15]. To accurately reconstruct their complex evolutionary history, it is therefore essential to identify the non-recombinant regions (NRRs), representing a single evolutionary history among the analysed genomes. We would expect some NRRs to have older and some younger origin times compared to estimates based on the entire genome, reflecting the complex patterns of recombination within and between different clades of lentiviruses over time.

In this study, we applied a mechanistic evolutionary model that takes into account the effects of recombination [15] and time-dependent changes in evolutionary rates [16] that acknowledges the contribution of epistatic changes at two or more sites in its calculation of a composite substitution rate. This was applied to reconstruct the evolutionary history of primate lentiviruses and to estimate the age of SIV lineages that gave rise to HIV-1 and HIV-2. We also estimated the time to the most recent common ancestor of SIV and the pSIV lentiviruses infecting lemurs and explored how this evolutionary reconstruction may explain the origins of primate lentiviruses.

## Methods

### Dataset

We followed the approach outlined by Bell and Bedford [15] to compile an initial dataset of 426 sequences from 24 host species. This dataset includes 5 to 31 sequences per host, incorporating all representative sequences from the Los Alamos National Laboratory (LANL) Compendium (http://www.hiv.lanl.gov/content/index) for each known SIV lineage, as well as each human-to-human transmissible HIV-1 and HIV-2 group. Specifically, we included 5 samples from HIV-1 group N (YBF30, DJO0131, SJGddd, U14296, and U14842), 8 samples from HIV-1 group O (96CMA102, ANT70, I_2478B, 99SE-MP1299, VAU, MVP5180, BCF06, and 99USTWLA), 3 samples from HIV-2 group A (BEN, ALI, and ST_JSP4), and 2 samples from HIV-2 group B (EHO and D205_ALT).

For HIV-1 group M, we included a total of 5 samples, one per subtype, excluding subtypes G, H, J, K, and recombinant subtypes to reduce the possible confounding effect of recombination (subtype G), and the low number of representative genomes (subtype H, J, and K) [17,18]. These 5 samples are isolates U455 (subtype A), WEAU160 (subtype B), ETH2220 (subtype C), 84ZR085 (subtype D), and VI850 (subtype F).

Additionally, we included 7 sequences of SIV collected from Bioko Island, each covering 472 to 633 bps from the conserved region of *pol* gene [12]. We also incorporated two reference samples of SIVdrl from LANL, isolates D3 and FAO, to estimate the divergence time between SIVdrl in Bioko and the mainland. Finally, we added a sequence from an endogenous lentivirus named grey mouse lemur prosimian immunodeficiency virus (pSIVgml), collected from conserved regions of *gag* and *pol* genes [10]. These 10 additional sequences were aligned using L-INSI algorithm implemented in mafft v7.520 [19], keeping the original 426 aligned sequences from ref [15] as a fixed template. The final dataset consisted of 436 aligned sequences from 28 host species.

### Comparison of recombination breakpoints

The inclusion of the 10 additional sequences did not alter the original 13 recombination breakpoints identified in ref [15] using a phylogenetic model in the HyPhy package GARD [20]. We further refined our dataset to include only sequences with 100 or more ungapped bases in each NRR. The final dataset consisted of 145 sequences for region 1 (2317 bps), 142 for region 2 (670 bps), 113 for region 3 (606 bps), 115 for region 4 (1295 bps), 237 for region 5 (936 bps), 246 for region 6 (1650 bps), 100 for region 7 (662 bps), 102 for region 8 (948 bps), 134 for region 9 (650 bps), 189 for region 10 (982 bps), 178 for region 11 (1050 bps), and 122 for region 12 (1826 bps) – see genome coordinates for each NRR in **Supplementary Figure 1**. The pSIVgml alignment spans NRRs 2 to 6, while the Bioko SIVblc, SIVprg, and SIVreg samples cover NRRs 5 and 6. The two SIVdrl samples from Bioko only cover part of NRR 6.

### Molecular clock inference

We initially assessed the temporal signal in our dataset using TempEst [21], observing no strong clock signal (**Supplementary Figure 2a**). Given the broadly similar molecular clock rates between various SIV and HIV lineages [9,22], we used HIV-1 group M to calibrate the molecular clock. We downloaded 100 complete-genome sequences of HIV-1 group M from LANL, with 20 samples from each subtype A, B, C, D, and F, aligning them with our main dataset as the template. After establishing the presence of a robust clock signal in this alignment (**Supplementary Figure 2b**), we divided the alignment into the 12 NRRs we identified earlier and separately inferred a clock rate for each region using an HKY+G4 substitution model, a strict clock with an uninformative CTMC rate prior [23] and constant population size coalescent prior in BEAST v.1.10 [24].

We then used these region-specific clock rates as priors for our dataset, using a normally distributed substitution rate prior with a mean that is equal to the HIV rates and a standard deviation that is 2.5 times wider than the posterior rate of HIV per NRR (see **Supplementary Figure 3** and **Supplementary Table 1**). The inferred posterior rate distributions from this analysis were used as the short-term substitution rate of SIV and HIV lineages in the PoW model. Finally, we evaluated temporal signals for each NRR in our dataset using BETS [25]. To estimate (log) marginal likelihoods, we used the generalised stepping-stone sampling method [26].

We ran all MCMC chains for 500 million generations, subsampling every 1 million iterations, and discarded the first 10% of the chains as burn-in. The log marginal-likelihood estimation comprised 100 path steps distributed according to quantiles from a β distribution with α=0.3, with each of the 101 resulting power posterior inferences running for 5×10^5^ iterations. We assessed convergence and mixing in Tracer v1.7.1 [27], ensuring that all relevant effective sample size values were >200.

### Reconstructing time trees using PoW model

To investigate the evolutionary timelines of HIV, SIV, and pSIV, we applied the PoW model [16]. This model accounts for the time-dependent decline in substitution rate estimates of viruses over time. Initially, we generated an ultrametric distance tree for each NRR with positive evidence for temporal signal (**Table 1**) using the HKY substitution model and a strict molecular clock.

**Table 1:**
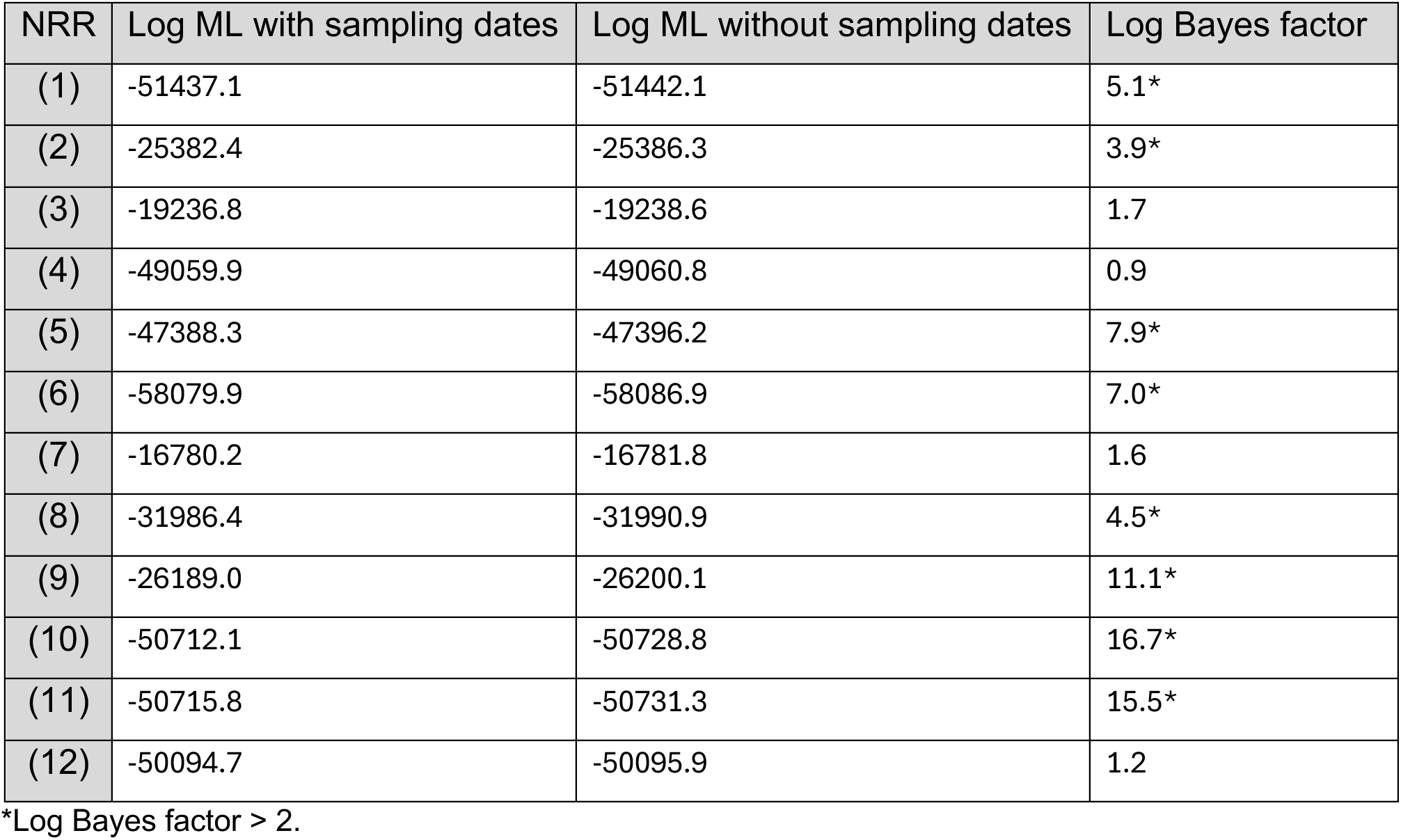
Evaluating the temporal signal present in the genomic samples for each non-recombinant region (NRR). We estimated log marginal likelihoods (ML) for each NRR using two datasets: (1) with sampling dates associated with the genomic samples and (2) a replicate dataset without the associated sampling dates.

| NRR | Log ML with sampling dates | Log ML without sampling dates | Log Bayes factor |
| --- | --- | --- | --- |
| (1) | -51437.1 | -51442.1 | 5.1* |
| (2) | -25382.4 | -25386.3 | 3.9* |
| (3) | -19236.8 | -19238.6 | 1.7 |
| (4) | -49059.9 | -49060.8 | 0.9 |
| (5) | -47388.3 | -47396.2 | 7.9* |
| (6) | -58079.9 | -58086.9 | 7.0* |
| (7) | -16780.2 | -16781.8 | 1.6 |
| (8) | -31986.4 | -31990.9 | 4.5* |
| (9) | -26189.0 | -26200.1 | 11.1* |
| (10) | -50712.1 | -50728.8 | 16.7* |
| (11) | -50715.8 | -50731.3 | 15.5* |
| (12) | -50094.7 | -50095.9 | 1.2 |

\*Log Bayes factor > 2.

The PoW model requires two parameters to transform ultrametric distance trees into time trees: the short-term substitution rate of SIV and HIV lineages and the substitution rate at the fastest-evolving rate group (μ_max_). The short-term substitution rate, inferred from the previous step, is approximately 1-2x10^-3^ s/s/y for most NRRs, except NRR 9 and 10 which had a nearly 2.25 and 1.7 times higher substitution rates (see **Supplementary Figure 3**). The μ_max_, set at 3.65x10^-2^ s/s/y in the model for all RNA viruses, was rescaled for NRRs 9 and 10 to account for their higher short-term substitution rates (8.21x10^-2^ s/s/y and 6.21x10^-2^ s/s/y, respectively). This parametrisation ensures that the proportion of sites evolving at various rates, from the fastest to the slowest (set to the substitution rate of 10^-9^ s/s/y), is accurately represented in the PoW model.

For constructing the time trees, we sampled 100 iterations from the post-burn-in posterior rate distributions and ultrametric distance trees for each NRR, excluding the initial 10% of MCMC chain runs. Each chain was run for 500 million generations, sampling every 1 million iterations. Finally, we used TreeAnnotator v.1.10.4 to merge the 100 reconstructed time trees into a maximum clade credibility tree, representing the most probable evolutionary history according to the PoW model.

### Incorporating the pSIVgml sample

Incorporating the genome sequence of pSIVgml —derived from a germline insertion approximately 4.2 million years ago [11] — into the PoW model for time tree construction requires an adjustment for the additional genetic divergence the sequence would have accumulated between its integration and the present day. This adjustment assumes a time-dependent evolutionary rate comparable to that observed in extant SIV and HIV lineages and is necessary to accurately estimate the total divergence since the MRCA of pSIVgml and contemporary SIV lineages within the PoW model framework.

To address this, we first estimated the expected genetic divergence for pSIVgml over 4.2 million years. This divergence was then apportioned between the branch leading to the pSIVgml node and the MRCA of SIV. To maintain ultrametricity in the distance trees used for downstream PoW model analysis, we distributed this divergence evenly between the pSIVgml branch and the stem branch leading to the SIV MRCA. These rescaled genetic distances were then converted into a time tree following the PoW procedure described above.

## Results

We prepared a dataset of 436 aligned sequences of human, simian, and prosimian immunodeficiency viruses from 28 hosts, including 26 nonhuman primate hosts for SIV, every human-to-human transmissible lineage of HIV (HIV-1 groups M, N, and O and HIV-2 groups A and B), and the sequence of the endogenous lentivirus, pSIVgml. We confirmed the presence of 13 recombination breakpoints within this dataset using GARD [15,20] and evaluated the strength of temporal signal in each of the 12 NRRs using BETS [25] (see **Methods**). We found positive evidence (log Bayes factor >2) for temporal clock signal in 8 NRRs (see **Table 1**). The clock rate in most NRRs varied between 1 to 2 x10^-3^ s/s/y, except for two NRRs in *env*, covering the gp120 subunit, which evolves at a rate approximately twice as high as the other regions (see **Supplementary Table 1**).

### Cross-validation of the PoW model with known divergence times

#### Dating human-to-human transmissible lineages of HIV

We reconstructed the evolutionary histories of each NRR with positive evidence for temporal signal using the PoW model (see **Methods**). We first validated the accuracy of the PoW over shallow timescales by re-estimating the tMRCAs of HIV-1 groups M, N, O and HIV-2 groups A and B (**Table 2a**). Our results confirmed previous findings that tMRCA estimates based on the *gag* and *env* loci are largely compatible with known estimates for the origins of these HIV clades, but that estimates based on the *pol* locus were consistently older [9,28].

**Table 2:** Estimates for time to the most recent common ancestor (tMRCA) of a selected set of HIV and SIV clades using the PoW model. **(a)** tMRCA estimates for HIV-1 groups M, N, O, and HIV-2 groups A, B in units of calendar time. **(b)** tMRCA estimates for SIV in sooty mangabeys (SIVsmm), chimpanzees (SIVcpz), SIV root, and divergence time between the SIV clade and the endogenous pSIVgml sample, shown in units of years ago (ya). All tMRCA estimates are shown for non-recombination regions (NRRs) 2 (part of *gag*), 6 (part of *pol*), and 11 (part of *env*). The pSIVgml genomic sample does not cover NRR 11. We also provide tMRCA estimates for the origin of HIV clades based on the genome-wide estimates. Dates in brackets indicate the 95% highest posterior density.

**(a)**
| Clade | *gag* (NRR 2) | *pol* (NRR 6) | *env* (NRR 11) | genome-wide |
| --- | --- | --- | --- | --- |
| HIV-1 M | 1890 (1740–1952) | 1822 (1513–1909) | 1912 (1845–1949) | 1884 (1857–1914) |
| HIV-1 N | 1961 (1922–1985) | 1976 (1938–1988) | 1986 (1975–1993) | 1972 (1963–1979) |
| HIV-1 O | 1888 (1758–1950) | 1855 (1623–1923) | 1874 (1788–1930) | 1897 (1857–1926) |
| HIV-2 A | 1934 (1802–1961) | 1913 (1798–1950) | 1894 (1771–1940) | 1941 (1924–1961) |
| HIV-2 B | 1954 (1903–1977) | 1939 (1836–1963) | 1951 (1919–1972) | 1946 (1930–1961) |

**(b)**
| Clade | *gag* (NRR 2) | *pol* (NRR 6) | *env* (NRR 11) |
| --- | --- | --- | --- |
| SIVsmm | 2.8 (1.2–8.9) Kya | 7.3 (3.0–34.6) Kya | 0.7 (0.3–1.5) Kya |
| SIVcpz | 27.5 (7.2–90.5) Kya | 21.5 (8.1–105.4) Kya | 25.8 (9.0–72.2) Kya |
| SIVroot | 0.3 (0.06–0.7) Mya | 1.8 (0.5–4.4) Mya | 0.8 (0.3–2.0) Mya |
| SIV/pSIV split | 16.9 (10.4–24.9) Mya | 50.5 (27.9–81.8) Mya | n/a |

We then sought to strengthen the phylogenetic signal present in the data by conducting a genome-wide estimation of the tMRCA for these HIV clades and found that the results were comparable to existing estimates of the origins of HIV-1 groups M and O in the late nineteenth century [29,30] and of HIV-2 groups A and B in the 1940s [22] (see **Table 2a**).

#### Dating various lineages of SIV

To estimate the tMRCA of SIVs on Bioko Island and their mainland counterparts, we used SIV samples of Bioko monkeys from a previous study [12]. These included 2 samples of SIV in drills (SIVdrl), 2 from Preuss’s guenon (SIVprg), 3 from black colobus (SIVblc), and 15 from red-eared guenon (SIVreg). The samples, typically 620-630 base pairs (bps) in length, spanned two NRRs in the *pol* gene (NRRs 5 and 6). The shortest samples, around 480 bps in length, from SIVdrl only covered a portion of NRR 6.

Without relying on calibrations at internal nodes or prior knowledge of Bioko Island’s separation from the mainland approximately 10,000–12,000 years ago (ya) [12], the PoW model estimated divergence times between several SIV lineages infecting monkeys on the island and their mainland counterparts that predated this geographical split. This included SIVblc-bioko from SIV in colobus monkeys (SIVcol), SIVreg-bioko from SIV in mona monkeys (SIVmon), and SIVprg-bioko from SIV in L’Hoest monkeys (SIVlst), all of which were predicted to have diverged ∼10-36 Kya (**Figure 1**; also see **Supplementary Figure 1**). However, the tMRCA for SIVdrl-bioko/SIVdrl-mainland split was estimated to be more recent, at 1.5 Kya (95% HPD: 0.4-3.3 Kya). This may be attributed to the shorter genomic fragments available for SIVdrl-bioko samples and potential under-sampling of the full diversity of SIVdrl on both the island and mainland.

**Figure 1:**
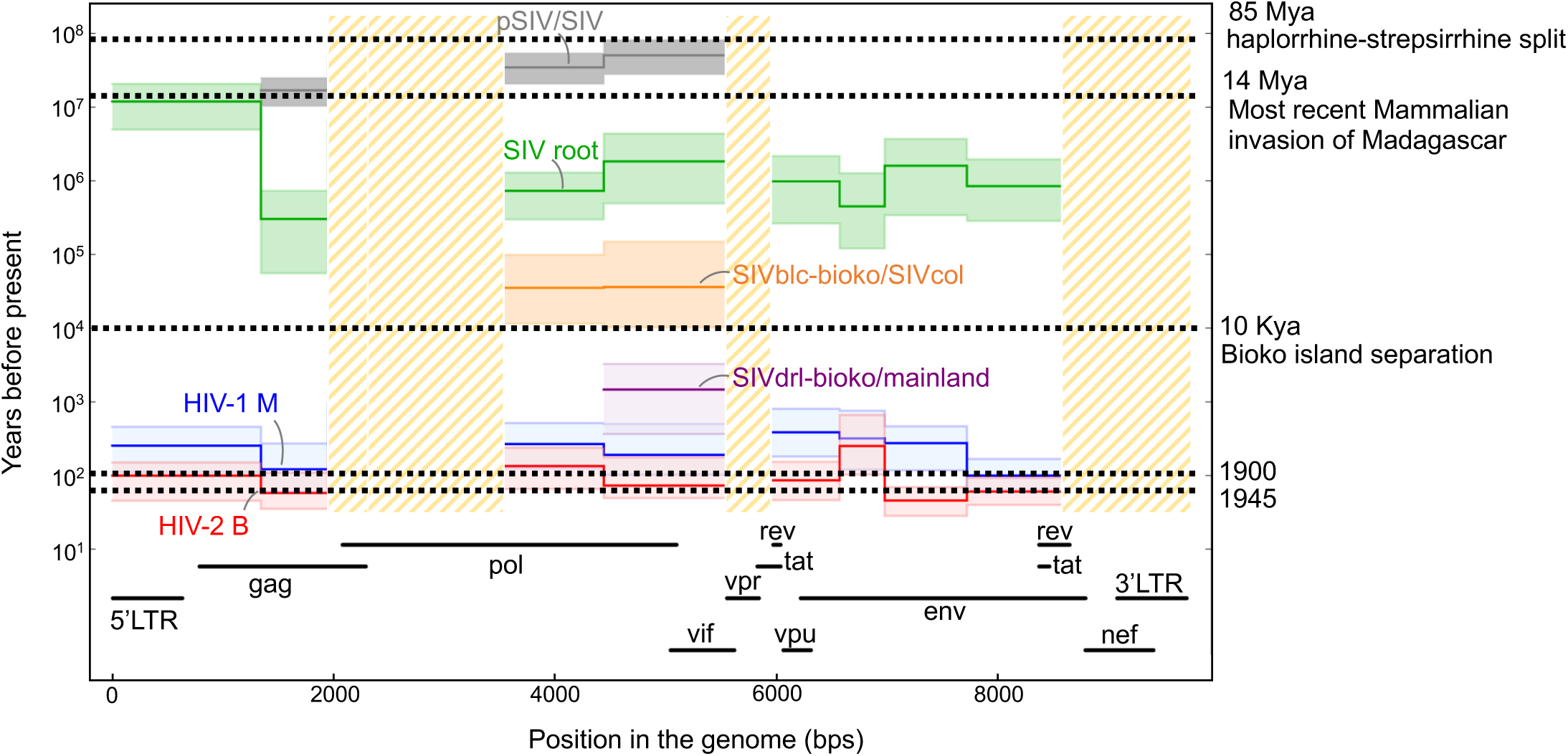
Time to the most recent common ancestor (tMRCA) estimates for primate lentiviruses using the PoW model. This timeline presents tMRCA estimates across various non-recombinant genomic regions for HIV-1 group M, HIV-2 group B, and the pSIV/SIV split, alongside SIV lineages from Bioko Island and the mainland, displayed on a logarithmic scale to represent years before present. The top of the timeline features the pSIV/SIV split, followed by the SIV root, with subsequent individual splits for SIV lineages from Bioko Island compared to their mainland counterparts depicted below. The lower part of the timeline shows the tMRCA for the HIV-1 M and HIV-2 B groups. Genomic regions are indicated below the timeline, with long terminal repeats (5’LTR and 3’LTR) and genes (*gag*, *pol*, *vif*, *vpr*, *vpu*, *rev*, *env*, *tat*, *nef*) detailed according to the HXB2 reference genome. Hashed areas represent genomic regions excluded from analysis due to the absence of a temporal signal, and error bars indicate the 95% highest posterior density (HPD) intervals for each estimate, with a solid line representing the median estimate. Species abbreviations for SIV lineages include drill (drl) and black colobus (blc).

Our estimates for the age of the SIV root in nearly all NRRs range from 300 thousand years (NRR 2, which covers part of *gag*) to 2 million years (NRR 6, which covers part of *pol*) before present (**Figure 1**). These findings are largely compatible with another study that used the separation of Bioko Island for clock calibrations with a relaxed molecular clock model to estimate the tMRCA of SIV [31]. These results further validate the reliability of the PoW model in estimating deep evolutionary events spanning millions of years, while also accurately recovering recent divergence times —on the order of hundreds of years— for the origins of various HIV groups, all within a unified evolutionary framework (see **Table 1** and **Supplementary Figure 4**).

### Evolutionary origins of SIV lineages that gave rise to HIV in humans

Our phylogenetic analysis showed SIVcpz, with a root age of between 30,000 to 200,000 years ago, is consistently older than SIVsmm, whose root age ranges from 300 to 11,000 years before present across all NRRs (**Figure 1**). These tMRCA estimates for both SIVcpz and SIVsmm, based on the PoW model, are up to several orders of magnitude older than those predicted by standard molecular clock methods that use calibrations based on contemporary sequences of SIV (**Supplementary Figure 4**).

One implication of the deep evolutionary origins of SIVcpz and SIVsmm is that humans may have been exposed to these viruses much longer than previously thought, since the MRCA of SIVcpz and SIVsmm and the MRCA of the human-to-human transmissible HIV lineages could be thousands to tens of thousands of years old (**Supplementary Figure 4**).

### Dating recent and deep virus-host associations

Our recombination-aware phylogenetic reconstruction also revealed the complex evolutionary history of SIVs (**Figure 2**; also see **Supplementary Figure 1**). In the majority of NRRs, SIV in colobus monkeys (SIVcol) consistently emerges as the outgroup, supported with strong posterior support. This observation is consistent with previous studies that suggest a long-term evolutionary isolation of SIVcol [15]. Our phylogenetic analysis suggests that SIVcol may have been coevolving with colobus monkeys for millions of years. Notably in NRR 1, which includes the 5’LTR and a portion of the *gag* gene, SIVcol appears as an outgroup with a very deep evolutionary branch of nearly 10 million years connecting it to the rest of the SIV lineages (**Supplementary Figure 1**).

**Figure 2:**
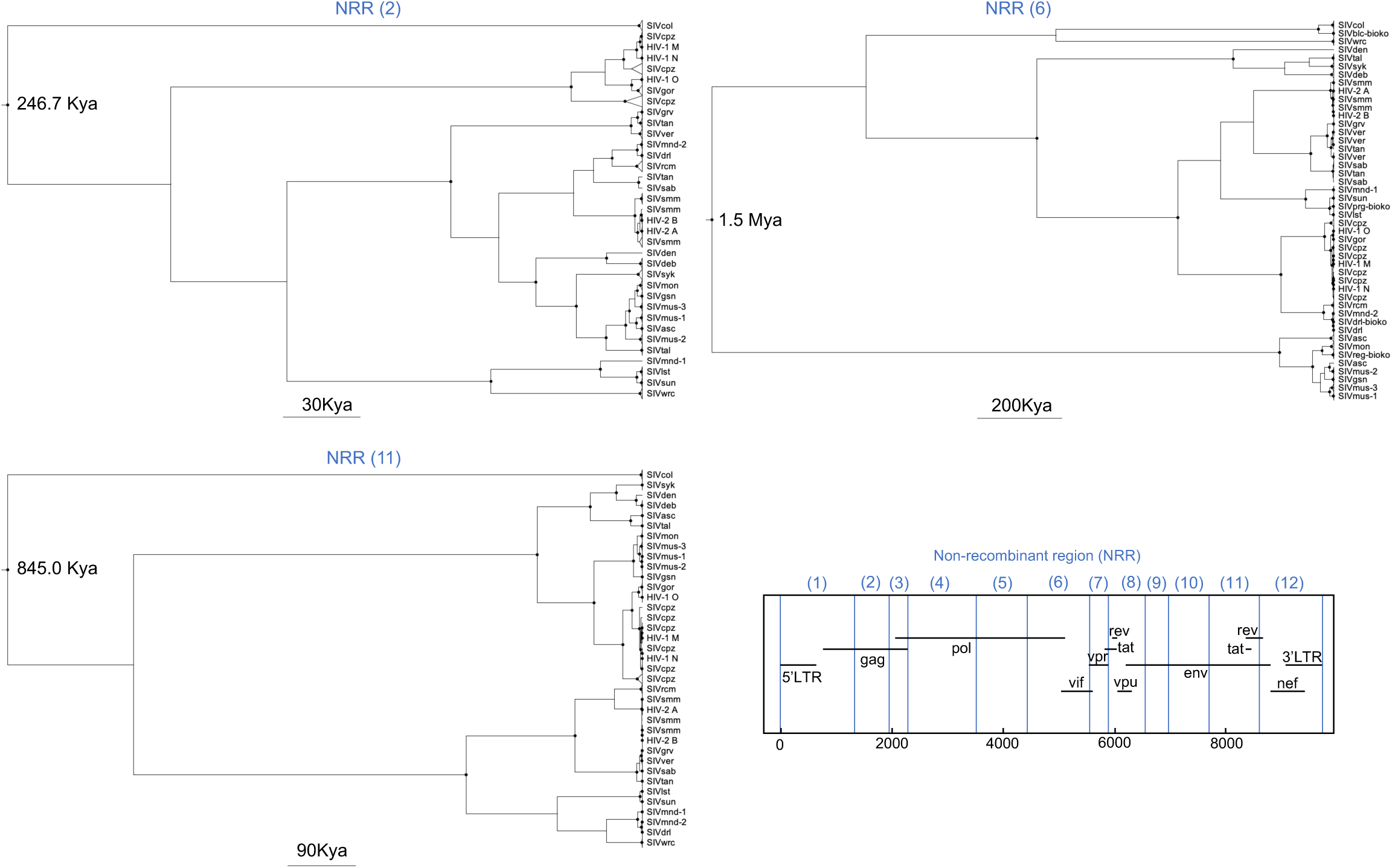
Maximum clade credibility time trees for a selected set of non-recombinant regions (NRRs) in primate lentivirus genome. This figure presents time trees for three key NRRs, specifically targeting the *gag* (NRR 2), *pol* (NRR 6), and *env* (NRR 11) loci. Each tree delineates the evolutionary trajectories of 141, 245, and 178 primate lentivirus samples, respectively, reconstructed using the Prisoner of War (PoW) model. Black circles show internal nodes with at least 90% posterior supports. The coordinates for each NRR are displayed at the bottom right, aligned with the HXB2 reference sequence. Species abbreviations for the SIV lineages include cpz (chimpanzee), gor (gorilla), mnd (mandrill), drl (drill), rcm (red-capped mangabey), sab (sabaeus monkey), tan (tantalus monkey), ver (vervet monkey), mal (malbrouck monkey), gri (grivet monkey), mac (rhesus macaque), mne (pig-tailed macaque), smm (sooty mangabey), lst (L’hoest’s monkey), sun (sun-tailed monkey), wrc (western red colobus), olc (olive colobus), col (colobus guereza), mus (moustached guenon), gsn (greater spot-nosed monkey), mon (mona monkey), asc (red-tailed guenon), tal (northern talapoin), deb (De Brazza’s monkey), den (Dent’s mona monkey), and syk (Sykes’s monkey).

We also found evidence supporting frequent host switching of SIV among African green monkey species. Our analysis corroborate previous findings [15] that in NRRs 2, 4, and 6, which encompass parts of the *gag* and *pol* loci, the SIVtan isolate, as detailed in [32], clusters with SIVsab and is distinctly separated from other SIVagm strains (**Supplementary Figure 1**). This suggests a relatively recent cross-species transmission of SIVsab from sabaeus to tantalus monkeys, likely within the last few centuries.

Consistent with other studies, we observed that SIVmnd-1 and SIVmnd-2 in mandrills do not consistently cluster together on the phylogenetic tree with strong posterior support, except in NRRs 11 (**Supplementary Figure 1**). This suggests that they may have shared a common ancestor anywhere from hundreds of thousands to millions of years ago. Similarly, SIVmus-1, SIVmus-2, and SIVmus-3 are identified as distinct lineages in most NRRs, likely sharing an MRCA dating back between 3,000 and 40,000 years (**Supplementary Figure 1**).

### Evolutionary origins of primate lentiviruses

The discovery of pSIV strains in strepsirrhine primates in Madagascar offers a unique perspective on the evolutionary timescales of primate lentiviruses. Although Madagascar parted from Africa around 160 Mya, primates only colonised the island around 65 Mya via rafting, island hopping, or temporary land bridges [33,34]. The presence of lentiviruses in Malagasy lemurs could have originated from several scenarios, as detailed in ref [10]:

1. host-virus cospeciation: This scenario dates back to the strepsirrhine-haplorrhine split, estimated at around 85 Mya.
2. Transfer between Malagasy Strepsirrhines and African Haplorrhines: This transfer could have occurred during the last mammalian colonisation of Madagascar, around14 Mya, possibly via a land bridge or a non-primate vector.
3. Aerial vector transfer: This scenario involves more recent, independent spillover events via aerial species such as bats.

While scenario 2 suggests a long-term association between lentivirus lineages and their hosts dating back to the latest terrestrial mammal invasion of Madagascar, scenario 3 proposes a possibly more recent and independent spillover events.

To assess which of these three scenarios are most likely, we used the PoW model to estimate the tMRCA of the SIV/pSIV split, factoring in the 4.2 Mya germline insertion of pSIVgml in *M. murinus* (see **Methods**). Our analysis, based on the *gag* gene (NRR 2), indicated a minimum MRCA age for SIV and pSIV of 17 Mya (95% HPD: 11–25 Mya), while the estimate based on the *pol* gene suggests a maximum age of 51 Mya (28–82 Mya). Both of these estimates are most consistent with the timing of the most recent primate colonisation of Madagascar and support scenario 2 as the most likely explanation for the origins of primate lentiviruses (see **Figure 1**). Neither of the two extreme tMRCA estimates align with the strepsirrhine-haplorrhine split around 85 Mya, thus ruling out host-virus cospeciation as proposed by scenario 1. Although scenario 3 remains plausible based on our results, it introduces additional complexity by requiring at least two independent spillover events.

## Discussion

Our findings demonstrated that the PoW model offers a robust and unified framework for reconstructing the evolutionary history of primate lentiviruses across a broad range of timescales. By explicitly accounting for time-dependent changes in viral evolutionary rates, the model overcomes key limitations of standard molecular clock approaches that rely on fixed shallow or deep calibration points. Notably, the PoW model accurately recovered the known origin times of the five human-to-human transmissible HIV strains (∼50–100 years ago), the divergence of SIV strains on Bioko Island (∼tens of thousands of years ago), and the root of the SIV lineage (∼millions of years ago)—all without the need for external biogeographic or palaeobiological calibrations.

We showed that SIV likely originated more than a million years ago and that the most recent common ancestor of many SIV lineages likely existed for hundreds of thousands of years in chimpanzees and colobus monkeys. More specifically, we found that SIVcpz, with a root age of between 30,000 to 200,000 years ago, is nearly ten times older than SIVsmm. This implies that human exposure to the SIV lineages in chimpanzees and sooty mangabeys that gave rise to HIV-1 and HIV-2, respectively, may have occurred for thousands of years before the pandemics began in the late 19th century (HIV-1) and mid-20th century (HIV-2), much longer than previously thought. The fact that such potential encounters never led to nascent HIVs becoming established in humans much earlier is likely due to changes in human social behaviour, increased population, mobility, and connectivity associated with urbanisation, and various other factors that have only emerged around or after the early 20^th^ century [7,35].

We also addressed one of the longstanding questions on the evolutionary origins of primate lentiviruses. By estimating the tMRCA of SIV and pSIV to be likely between 17 and 51 million years old, we were able to rule out ancestral codivergence of primate lentiviruses into haplorrhine and strepsirrhine lineages as a likely scenario of primate lentivirus origins. Instead, we found the land-bridge hypothesis, whereby a terrestrial mammal or a non-primate vector transferred lentiviruses between haplorrhine and strepsirrhine in mainland Africa and Madagascar, to be a more plausible scenario.

Our findings can also offer new insights into understanding the evolution of ancient innate immune defence mechanisms in simian and prosimian hosts against lentivirus infections. The tens to hundreds of thousands of years of codivergence between many SIV strains and their simian hosts, as estimated by the PoW model, may provide the opportunity for the host to develop effective immune defence mechanisms to tolerate infection through control immune activation [36]. Also, similarities in set point viral loads of minimally pathogenic SIVsmm in sooty mangabeys with those of highly pathogenic HIV-1 in humans (or indeed of SIVsmm in the Asian rhesus macaques) [37] may indeed reflect differences in degrees of virus-host adaptation and resistance to virus replication-induced pathology that reflect their distinct evolutionary histories and host associations [38].

While our tMRCA estimates for HIV-1 group M and HIV-2 groups A and B aligned broadly with those from standard molecular clock methods, the PoW model produced substantially older estimates for the origins of SIVcpz and SIVsmm. This discrepancy underscores a central limitation of conventional approaches: their failure to account for time-dependent rate variation results in systematic underestimation of divergence times beyond ∼100 years, primarily due to substitution saturation [39,40]. More broadly, our findings expose the inherent flaws in fixed-point calibration methods. Calibrations based on recent sequences cannot recover deep viral histories, while those based on biogeographic or palaeobiological events cannot resolve recent divergences. These approaches also distort timelines by assuming constant rates across timescales.

By contrast, the PoW model accommodates continuous rate decay, enabling consistent reconstruction of divergence events from decades to millions of years without relying on individual internal node calibrations. By integrating lentivirus data across recent, intermediate, and ancient timescales, our analysis reconciles previously fragmented perspectives on SIV evolution—spanning contemporary HIV epidemics, Late Pleistocene SIV divergence events (Bioko Island separation), and ancient lentiviral insertions (pSIVgml)—and reveals that while host–virus codivergence is limited, the deep antiquity of lentiviruses is strongly supported by multiple lines of evidence.

The PoW model was previously used to reconstruct the evolutionary origins of the sarbecovirus lineage [16,41] and was found to be in remarkable concordance with signatures of a selection of human genomic datasets that indicate an arms race with corona-like viruses dating back to 25,000 years BP [42], providing an external comparator for this methodology. Future research could apply this model more broadly to other viruses to re-evaluate their evolutionary timescales and gain insights into host– virus relationships that are critical for understanding viral pathogenicity, guiding antiviral development, and assessing pandemic risk—including estimating the timing of cross-species transmission events, zoonotic emergence, and host range expansion across diverse virus families.

## Supporting information

alignments and pow-transformed trees

## Data and code availability

All sequence alignments and PoW-transformed phylogenetic trees are provided in the supplementary materials.

## Acknowledgment

This work was supported by the Biotechnology and Biological Science Research Council (BB/M011224/1 to M.G.) and the European Research Council (101001623-PALVIREVOL to A.K.).

## Supplementary Figures

**Supplementary Figure 1:**
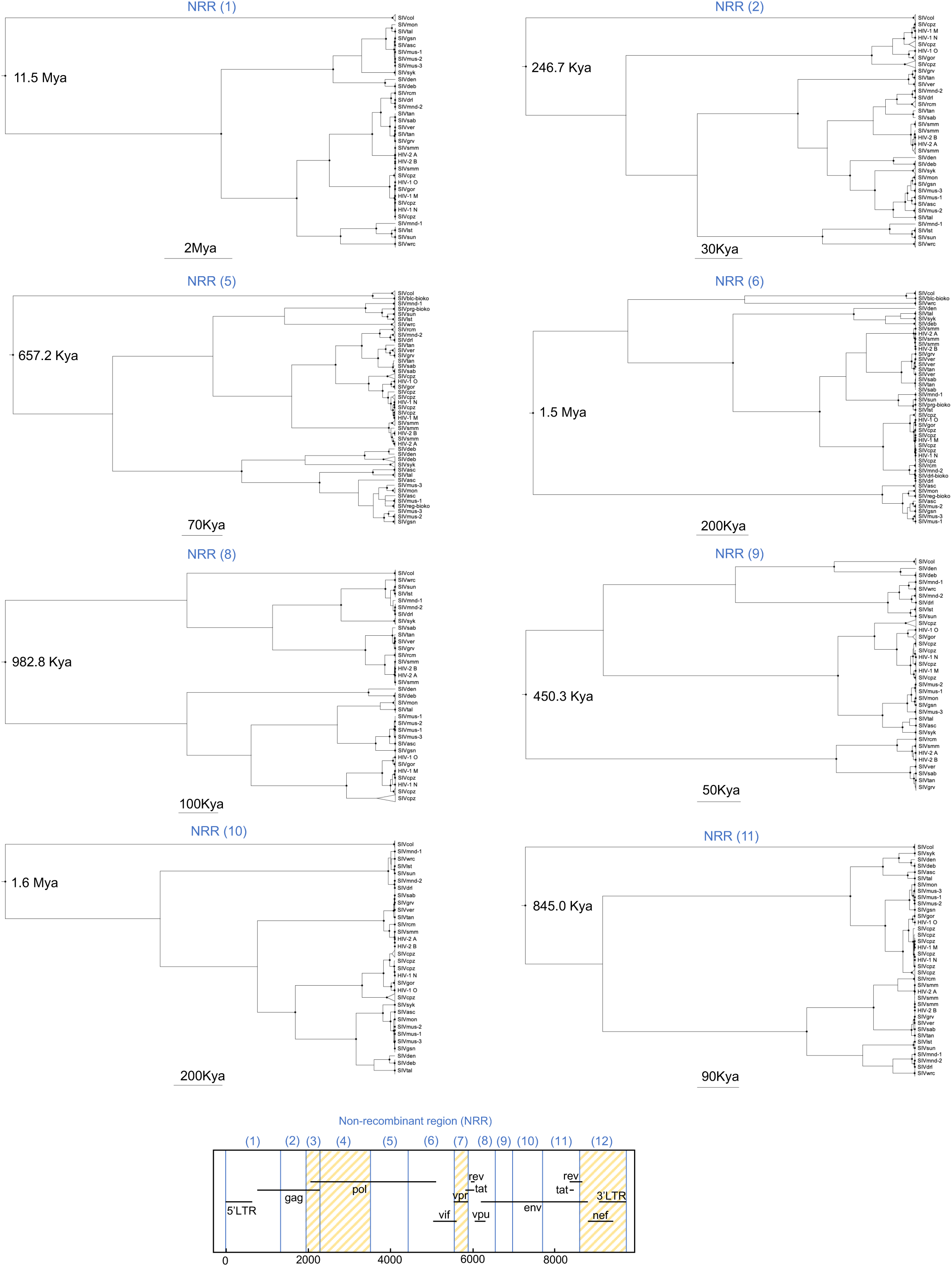
Maximum clade credibility time trees of primate lentiviruses across non-recombinant regions (NRRs) with positive evidence for temporal signal. This figure displays 10 time trees representing the evolutionary history of 435 primate lentivirus samples (excluding pSIVgml), reconstructed using the Prisoner of War (PoW) model for each NRR. These trees illustrate the temporal diversification patterns and provide insights into the lineage-specific evolutionary dynamics of SIV and HIV. For enhanced clarity, monophyletic clades from the same lineage are collapsed into triangles, the width of which is proportional to the number of sequences within. Most NRRs identify SIVcol as an outgroup, while SIVsmm is consistently clustered closely with HIV-2 groups A and B. Similarly, SIVcpz is clustered with various HIV-1 groups (M, O, and N), with SIVgor positioned as the immediate outgroup to HIV-1 group O. The pSIVgml sequence, only covering NRRs 2 to 6 and consistently appearing as an outgroup to all SIV lineages (not shown), is connected to the primate lentivirus root by a deep branch (>14 million years). The coordinates for each NRR are indicated at the bottom according to the HXB2 reference. Hashed areas represent genomic regions excluded from analysis due to the absence of a temporal signal. Black circles show internal nodes with at least 90% posterior supports. Species abbreviations for the SIV lineages are cpz (chimpanzee), gor (gorilla), mnd (mandrill), drl (drill), rcm (red-capped mangabey), sab (sabaeus monkey), tan (tantalus monkey), ver (vervet monkey), mal (malbrouck monkey), gri (grivet monkey), mac (*rhesus macaque*), mne (pig-tailed macaque), smm (sootey mangabey), lst (L’hoest’s monkey), sun (sun-tailed monkey), wrc (western red colobus), olc (olive colobus), col (*colobus guereza*), mus (moustached guenon), gsn (greater spot-nosed monkey), mon (mona monkey), asc (red-tailed guenon), tal (northern talapoin), deb (de brazza’s monkey), den (dent’s mona monkey), and syk (sykes’s monkey).

**Supplementary Figure 2:**
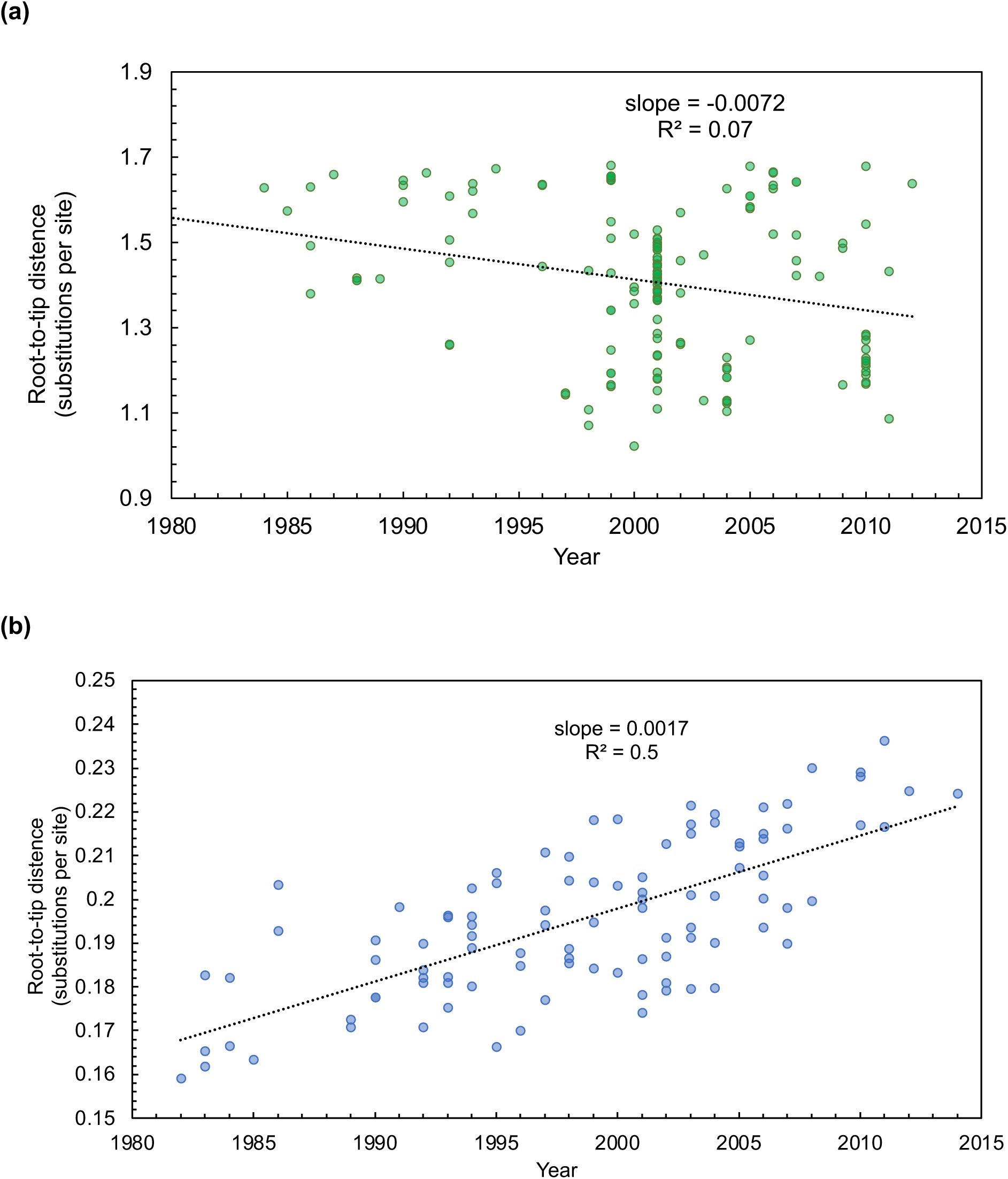
Evaluating temporal signal in the time-stamped data. Root-to-tip distance as a function of sampling time for **(a)** a merged dataset of SIV and HIV samples **(b)** HIV-1 group M samples. The plots are based on maximum likelihood tree reconstructions with a root position that maximises the residual mean squared for the regression of root-to-tip divergence and sampling time (black dotted line). In each panel, the regression line is shown in black dotted line along with the corresponding slope (evolutionary rate) and coefficient of determination, R^2^.

**Supplementary Figure 3:**
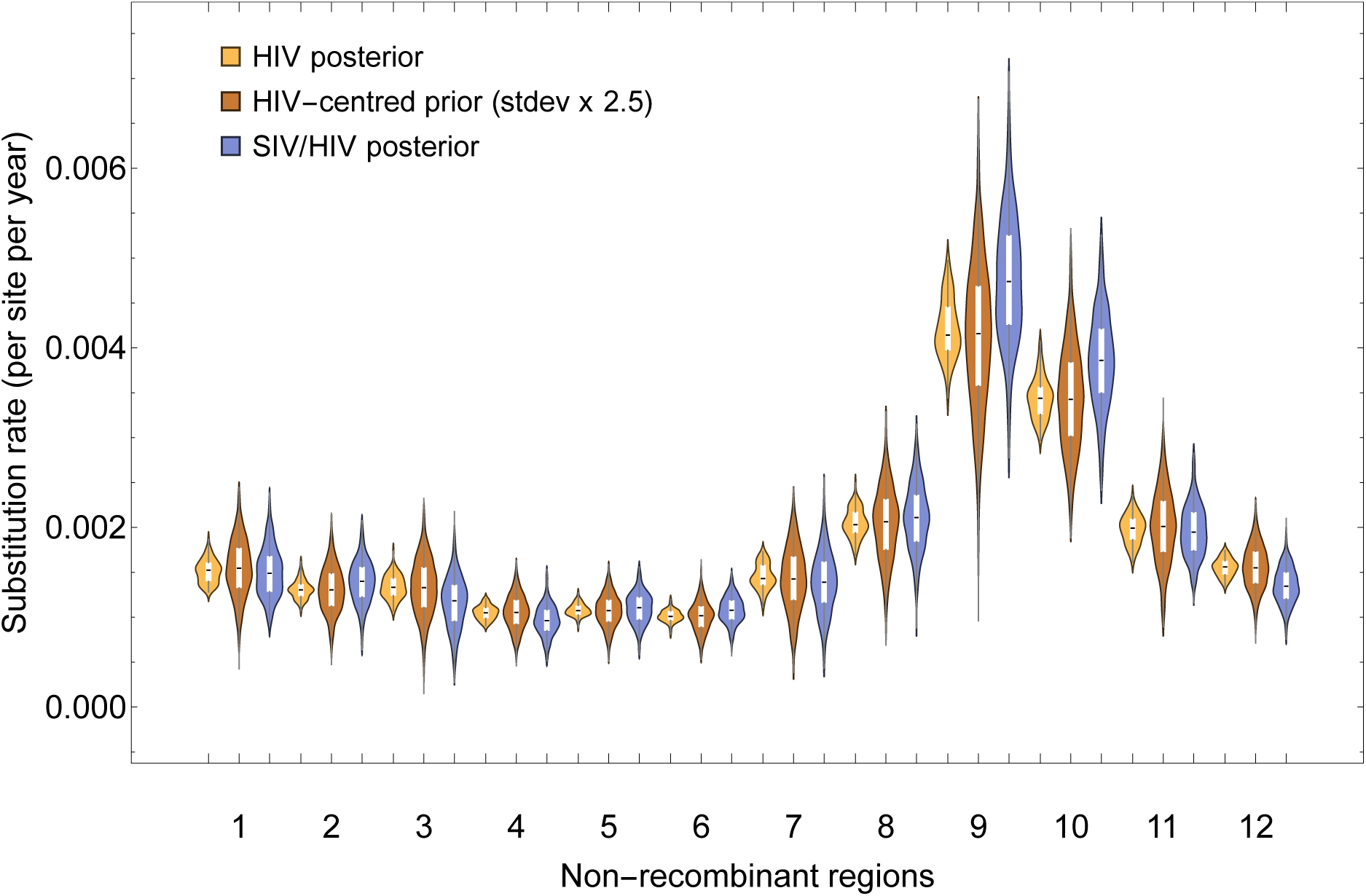
Substitution rates for the 12 non-recombinant regions (NRRs) of HIV and SIV samples. The inferred posterior distribution of substitution rates for HIV sequences (yellow), derived from an analysis of 100 HIV-1 group M time-stamped sequences, is used as the prior (red) to infer the substitution rate of the combined SIV and HIV dataset (blue). For each NRR, the rate prior is normally distributed with a median that is equal to the HIV dataset and a standard deviation that is 2.5 times higher than that of the HIV dataset. Each violin plot corresponds to a distinct NRR, numbered 1 through 12. The width of each plot reflects the probability density of rates estimated using a separate HKY+G4 substitution model and a relaxed uncorrelated lognormal molecular clock, implemented in BEAST v.1.10.

**Supplementary Figure 4:**
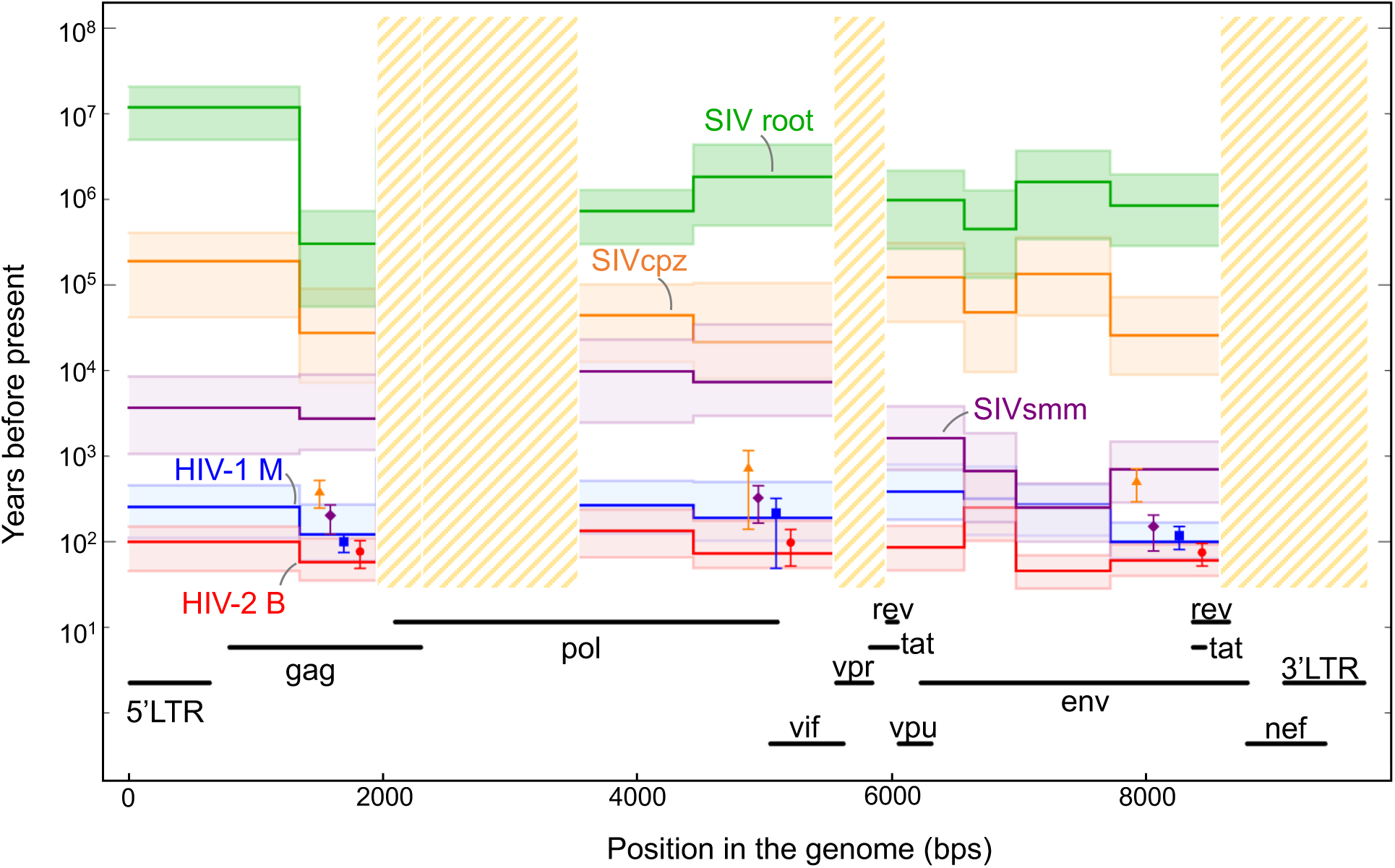
Time to the most recent common ancestor (tMRCA) estimates for SIV lineages leading to HIV-1 and HIV-2. This timeline shows tMRCA estimates across various non-recombinant regions with positive evidence of temporal signal for HIV-1 group M (in blue), HIV-2 group B (in red), and the roots of SIVcpz (in orange) and SIVsmm (in purple), with the SIV root age represented in green. Estimates are plotted along a logarithmic scale, indicating years before the present. Data points with error bars represent tMRCA estimates derived from a standard molecular clock method using a relaxed clock, located on the timeline at NRR 2, 6, and 11, based on an earlier study. While these standard molecular clock estimates generally concur with the PoW model for the age of HIV-1 and HIV-2 groups, the PoW model suggests substantially older estimates for SIVsmm and SIVcpz roots, indicating a much deeper evolutionary history for these lineages. Genomic regions are indicated below the timeline, with long terminal repeats (5’LTR and 3’LTR) and genes (*gag*, *pol*, *vif*, *vpr*, *vpu*, *rev*, *env*, *tat*, *nef*) detailed according to the HXB2 reference genome. Hashed areas represent genomic regions excluded from analysis due to the absence of a temporal signal, and error bars indicate the 95% highest posterior density (HPD) intervals for each estimate, with a solid line representing the median estimate. Species abbreviations for SIV lineages include chimpanzee (cpz) and sooty mangabey (smm).

## Supplementary Tables

**Supplementary Table 1:**
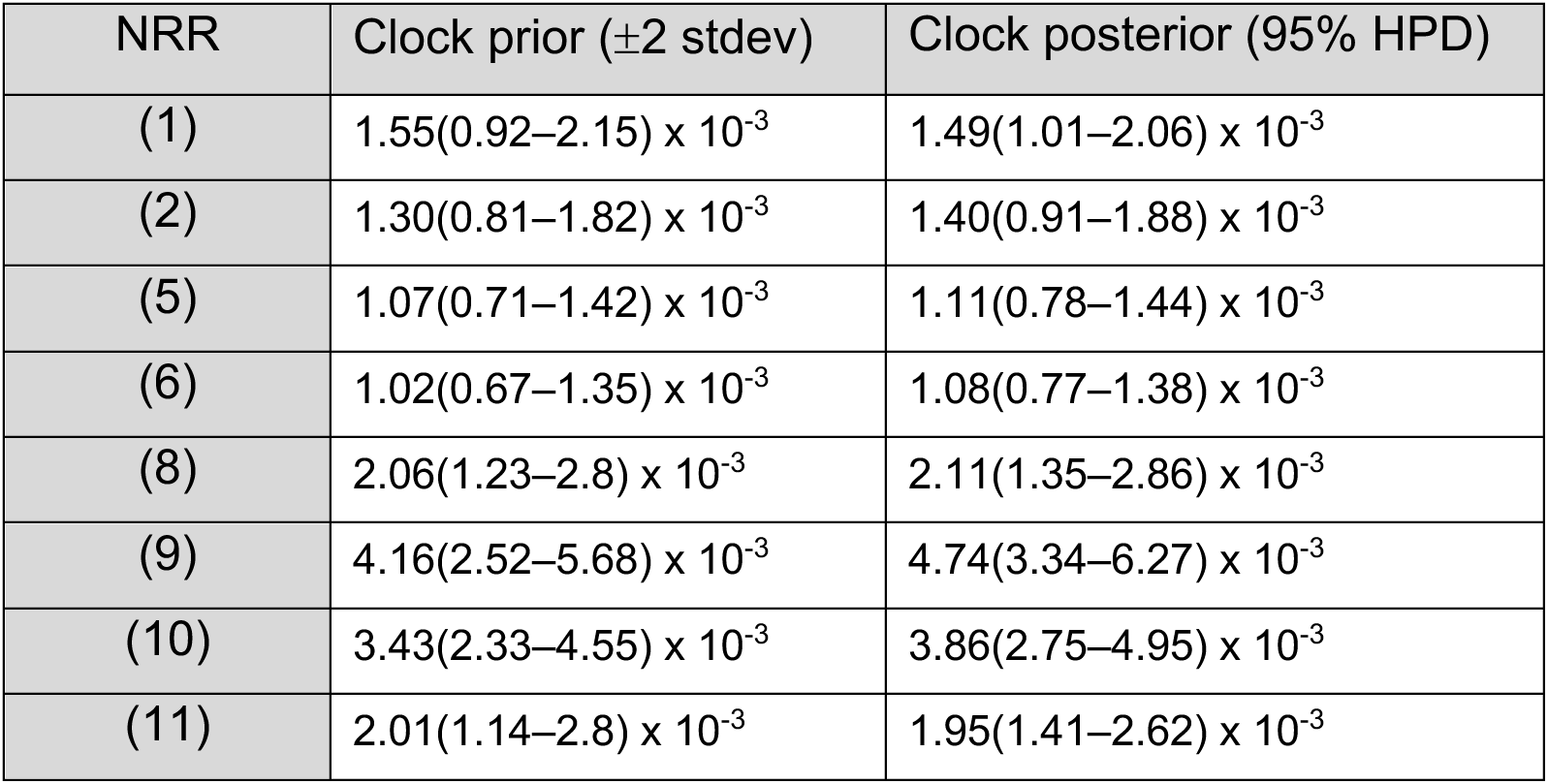
Clock calibration prior and posterior clock rates for each non-recombinant region (NRR) with positive evidence for temporal signal. Clock calibration priors per NRR are based on 100 HIV genomic samples over a 35-year timespan, with very strong clock signal. Clock rate posteriors are for the curated dataset of SIV and HIV genomic samples with known collection dates over the same 35-year timespan.

| NRR | Clock prior ($\pm 2$ stdev) | Clock posterior (95% HPD) |
| --- | --- | --- |
| (1) | $1.55(0.92-2.15) \times 10^{-3}$ | $1.49(1.01-2.06) \times 10^{-3}$ |
| (2) | $1.30(0.81-1.82) \times 10^{-3}$ | $1.40(0.91-1.88) \times 10^{-3}$ |
| (5) | $1.07(0.71-1.42) \times 10^{-3}$ | $1.11(0.78-1.44) \times 10^{-3}$ |
| (6) | $1.02(0.67-1.35) \times 10^{-3}$ | $1.08(0.77-1.38) \times 10^{-3}$ |
| (8) | $2.06(1.23-2.8) \times 10^{-3}$ | $2.11(1.35-2.86) \times 10^{-3}$ |
| (9) | $4.16(2.52-5.68) \times 10^{-3}$ | $4.74(3.34-6.27) \times 10^{-3}$ |
| (10) | $3.43(2.33-4.55) \times 10^{-3}$ | $3.86(2.75-4.95) \times 10^{-3}$ |
| (11) | $2.01(1.14-2.8) \times 10^{-3}$ | $1.95(1.41-2.62) \times 10^{-3}$ |

